# Acute ethanol stress induces sumoylation of conserved chromatin structural proteins in *Saccharomyces cerevisiae*

**DOI:** 10.1101/2020.11.09.375147

**Authors:** Amanda I. Bradley, Nicole M. Marsh, Heather R. Borror, Kaitlyn E. Mostoller, Amber I. Gama, Richard G. Gardner

## Abstract

Stress is a ubiquitous part of life that disrupts cellular function and, if unresolved, can irreparably damage essential biomolecules and organelles. All organisms can experience stress in the form of unfavorable environmental conditions including exposure to extreme temperatures, hypoxia, reactive oxygen species, alcohol, or shifts in osmolarity. To survive, organisms must sense these changes then react and adapt. One highly conserved adaptive response to stress is through protein sumoylation, which is a post-translational modification by the <u>s</u>mall <u>u</u>biquitin-like <u>mo</u>difier (SUMO) protein. In this study, we examine the effects of acute ethanol stress on protein sumoylation in the budding yeast *Saccharomyces cerevisiae*. Although ethanol induces protein sumoylation, the targets and roles of sumoylation are largely unknown. Here, we found that cells exhibit a transient sumoylation response after exposure of cells to ≤ 7.5% volume/volume ethanol. The response peaks at 15 minutes and resolves by 60 minutes, indicating that cells have an adaptive response to low concentrations of ethanol. By contrast, the sumoylation response becomes chronic at 10% ethanol stress with no resolution by 60 minutes. To identify key targets of ethanol-induced sumoylation, we performed mass spectrometry analyses and identified 18 proteins with increased sumoylation after acute ethanol exposure, with 15 identified as known chromatin-associated proteins. Two of these proteins are the chromatin structural proteins Smc5 and Smc6, which are sumoylated by the activity of the SUMO ligase Mms21. Ethanol-induced Smc5/6 sumoylation occurs during G1 and G2M phases of the cell cycle but is abrogated during S phase despite the fact that other proteins are sumoylated during this phase. Acute ethanol exposure leads to formation of Rad52 foci indicating DNA damage similar to that observed with the addition of methyl methanesulfonate (MMS), which is an alkylating agent that damages DNA. Whereas MMS exposure induces the intra-S phase DNA damage checkpoint as observed by Rad53 phosphorylation, ethanol exposure does not induce the intra-S phase checkpoint and prevents Rad53 phosphorylation when added with MMS. From these results, we propose that ethanol induces a structural change in chromatin, possibly through DNA damage, and this causes sumoylation of conserved chromatin-associated proteins, including Smc5 and Smc6.

## Introduction

Stress is an inevitable consequence of life. Every organism experiences this unwelcome and detrimental phenomenon. At the cellular level, stress is often caused by alterations in intra- or extra-cellular environments. Prolonged exposure of cells to stress conditions such as oxidation, temperature shifts, hypoxia, osmolarity alterations, genotoxic events, and a multitude of others can lead to damage of DNA, RNA, proteins, and other macromolecules. Consequently, the ability to sense and adapt to changing extracellular conditions is integral to cell survival. An effective response to exogenous stressors is elicited through activation of cellular stress response pathways that alter gene expression and/or protein interactions or activity in a coordinated effort to re-establish and maintain homeostasis (Galluzzi et al., 2018). The inability to respond quickly and adapt to stress can lead to cell death, and failure to adapt to prolonged stress conditions underlies many human pathologies such as heart disease, neurodegeneration, and cancer (Fulda et al., 2010).

Post-translational modifications (PTMs) play a key role in aiding cell survival during stress conditions. One PTM found to increase during a number of stress conditions is the <u>s</u>mall <u>u</u>biquitin like <u>mo</u>difier (SUMO). Similar to protein ubiquitination, protein sumoylation utilizes an enzymatic cascade that leads to attachment of SUMO molecules to target substrates (Johnson et al., 1997; Okuma et al., 1999). SUMO modifications can alter protein localization, protein-protein interactions, aid in protein stability and solubility (Geiss-Friedlander and Melchior, 2007). While protein sumoylation is known to be increased across a broad array of stresses, the majority of the target proteins and the kinetics by which they are sumoylated are distinct (Guo and Henley, 2014; Lewicki et al., 2015; Tempé et al., 2008). Although studies have reported the involvement of protein sumoylation in cellular stress responses and various targets (Miller et al., 2013; Oeser et al., 2016; Zhou et al., 2004), the function of specific protein sumoylation events during distinct stresses still remains poorly understood.

To better understand the key targets and functions of protein sumoylation during stress conditions, we have been recently studying the proteins that become sumoylated during acute ethanol exposure in the budding yeast *Saccharomyces cerevisiae*. The utilization of different yeasts for the purpose of ethanol production has been exploited for centuries (Mohd Azhar et al., 2017; Parapouli et al., 2020). Due to the industrial importance of ethanol production, a considerable amount of research has examined the differences in ethanol tolerance among laboratory and industrial yeast strains (Lewis et al., 2010; Steensels and Verstrepen, 2014). While yeast cells can tolerate relatively high concentrations of ethanol, this does not prevent them from experiencing cellular stress during acute and chronic exposure to ethanol. Chronic exposure to high concentrations of ethanol has been shown to alter membrane fluidity and lipid composition, increase reactive oxygen species (ROS) production through decoupling oxidative phosphorylation in the mitochondria, and cause protein misfolding (Auesukaree, 2017). In this study, we explore the effects of acute ethanol stress on protein sumoylation. From a mass spectrometry analysis, we found 18 proteins that appear be sumoylated upon ethanol acute exposure, 7 of which are transcription factors. We also found that the members of the chromatin structural Smc5/Smc6 complex are sumoylated upon acute exposure to ethanol. Overall, the major targets for ethanol-induced sumoylation are chromatin-associated proteins involved in transcriptional regulation or chromatin structure.

## Results

### Global sumoylation kinetics in yeast dependent on ethanol concentration

We previously examined global sumoylation response patterns over time in *S. cerevisiae* to various stressors that included ethanol (10% v/v) (Oeser et al., 2016). In that study, acute exposure to high ethanol stress resulted in a steady accumulation of SUMO conjugates over a 60-minute time course. While we were interested in sumoylation induced by 10% v/v ethanol, we also wanted to determine if the sumoylation patterns observed remained unchanged at ethanol concentrations lower that 10% (v/v), or if the sumoylation effect was only observed at ethanol concentrations that limit yeast growth. Utilizing a yeast strain where the endogenous SUMO gene, *SMT3*, was tagged with a His_6_-FLAG sequence at its 5’ end (Fig 1A), we examined sumoylation during ethanol stress at the following concentrations (v/v): 1%, 2.5%, 5%, 7.5%, and 10%. At concentrations lower than 10%, we found that the ethanol sumoylation response becomes transient with a pronounced increase in SUMO conjugates at 15 minutes that returned to basal levels by 60 minutes (Fig 1B), indicating that at lower ethanol concentrations yeast can mount an adaptive response that mitigates the need for chronic sumoylation.

**Figure 1.**
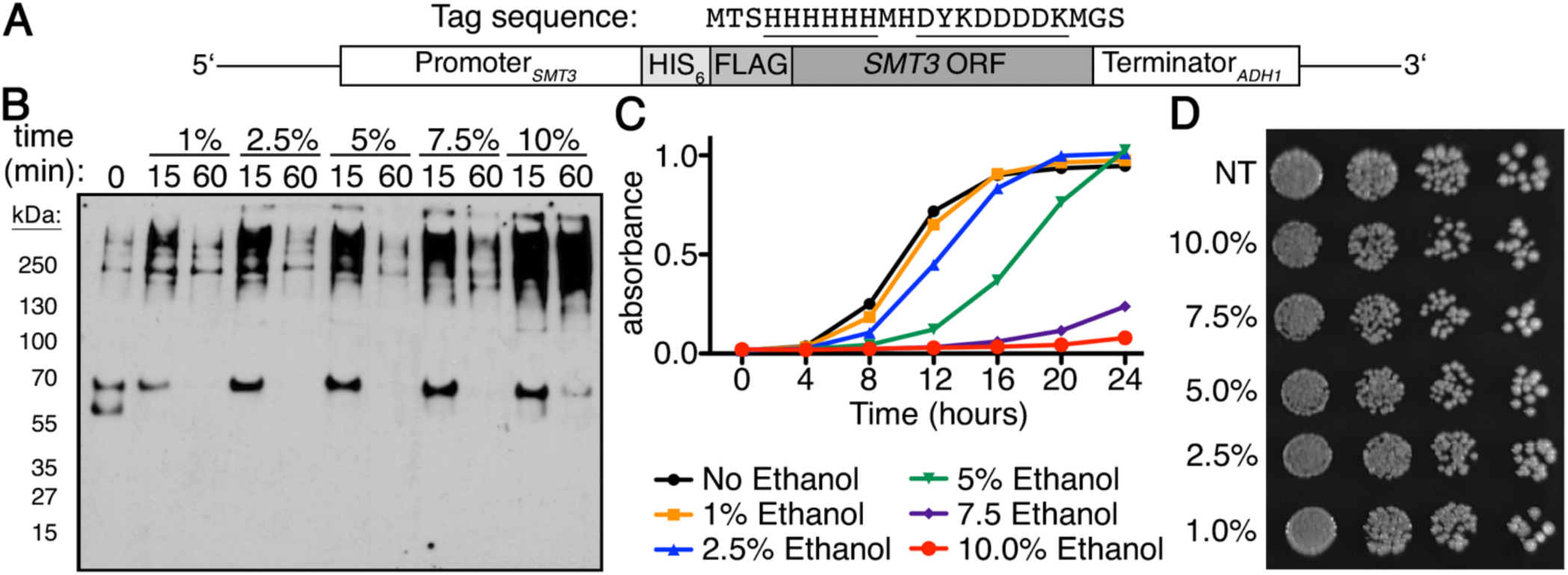
Global sumoylation kinetics are dependent on ethanol concentration. (A) Schematic for His_6-_FLAG-SMT3 located at the *SMT3* locus and expressed from the endogenous *SMT3* promoter. His_6_-FLAG sequence shown, sequences underlined. (B) Comparison of global sumoylation changes that occur during cellular exposure of ethanol (1%, 2.5%, 5%, 7.5%, and 10% v/v) over 60 minutes. Changes in sumoylation patterns were examined by Western analysis using an anti-FLAG antibody. (C) Quantitative measure of growth rates in liquid culture generated by Bioscreen C automated growth curve analysis. Cells were grown in triplicate at 30°C in rich media with 0%, 1%, 2.5%, 5%, 7.5%, or 10% ethanol for 24 hours with continuous shaking. Absorbance at 600nm was measured every 30 minutes and average absorbance (at 600nm) was plotted versus time. Error bars show SD for triplicate samples. (D) Growth in ethanol does not affect viability. His_6_-FLAG-*SMT3* cells were grown to mid-log phase in rich liquid media then treated for 60 minutes in rich media with 0%, 1%, 2.5%, 5%, 7.5%, or 10% ethanol before being 10-fold serially diluted onto rich media plates and incubated at 30°C for 3 days.

To determine what levels of ethanol affect yeast cell growth, we queried the effect of ethanol concentration on cell growth when cells are chronically exposed to each of these concentrations. To do this, we performed liquid culture growth assays over a period of 24 hours. Cells in 1% ethanol exhibited growth identical to cells with no ethanol treatment, while cells in 2.5% and 5% ethanol were delayed before entering into exponential growth. Cells in 7.5% and 10% ethanol, however, did not achieve exponential growth during the 24-hour period (Fig 1C). To be certain that acute exposure to ethanol did not impact cell viability, we performed spot dilution tests on media lacking ethanol after the cells were exposed to the same concentrations of ethanol as listed above for 1 hour in liquid culture. We did not observe any growth deficiency between untreated and ethanol-treated cells (Fig 1D), indicating that acute ethanol exposure does not alter cell viability as observed with chronic exposure (Fig 1C).

### A chromatin structure complex is sumoylated during acute ethanol exposure

To identify proteins sumoylated during acute ethanol stress, we utilized a label-free mass spectrometry (MS) approach as previously described (Oeser et al., 2016). Although we found that sumoylation is transient at lower ethanol concentrations, we chose to query sumoylation at 10% ethanol to maximize the potential proteins that could be identified at early and late time points. Similar to our previous studies (Oeser et al., 2016), we enriched for sumoylated species from His_6_-FLAG-*SMT3* cell lysates derived from cultures prior to ethanol treatment and after ethanol treatment, in this case 5 and 60 minutes post-ethanol exposure (Fig 2A). Sumoylated proteins were isolated by gel purification and subject to trypsin digestion prior to MS analysis. All peptide counts were examined to determine which proteins showed increases at 5 minutes or 60 minutes of 10% ethanol exposure.

**Figure 2.**
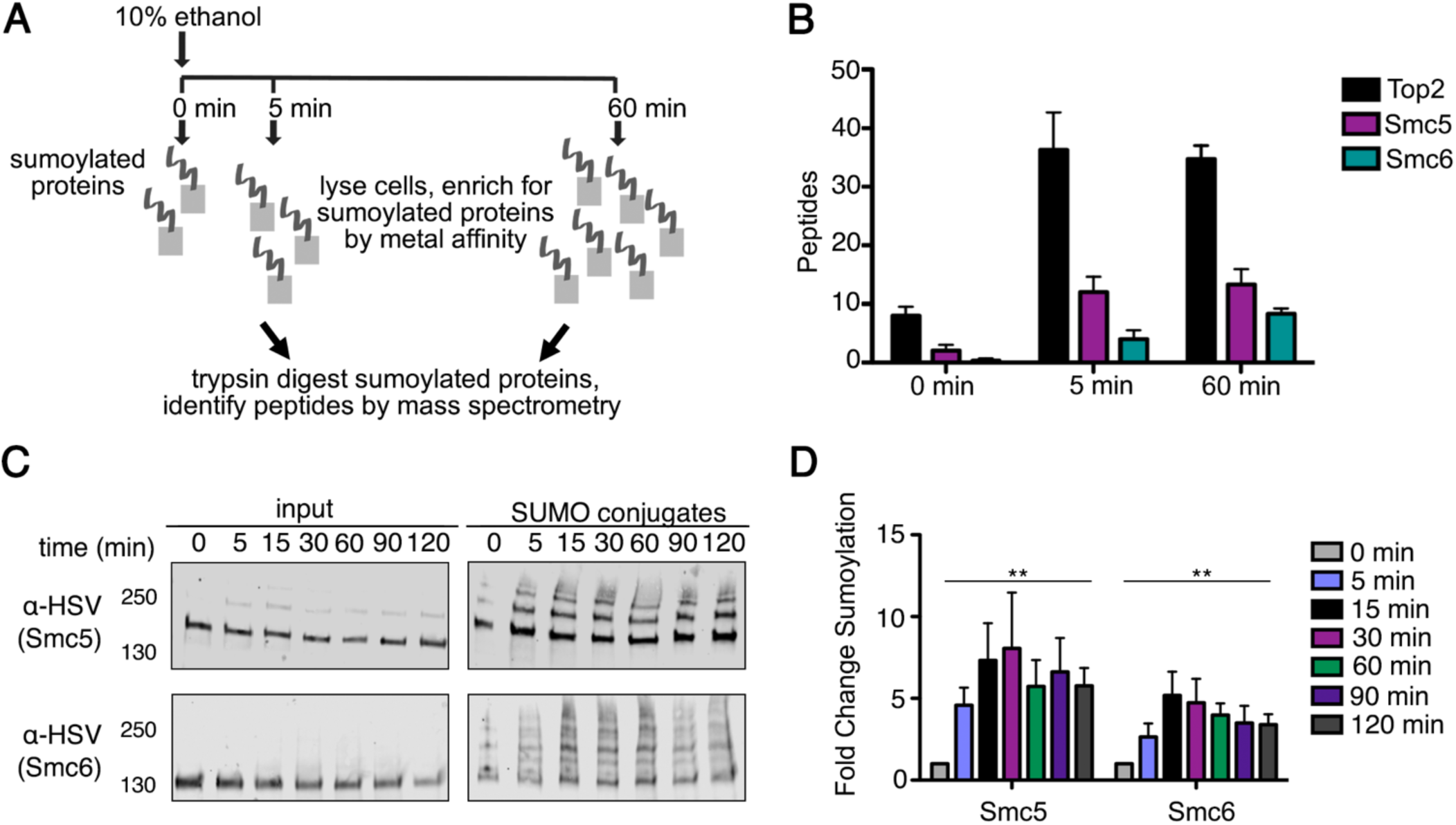
The Smc5/Smc6 chromatin complex is sumoylated during acute ethanol exposure. (A) Mass spectrometry strategy to identify proteins sumoylated during acute ethanol exposure. (B) Total peptide counts identified for the top 3 proteins (Top2, Smc5, and Smc6) at 0, 5 and 60 minutes of 10% ethanol stress. (C) Cells expressing His_6_-FLAG-*SMT3* and either 3xHSV epitope tagged Smc5 or Smc6 from their endogenous promoters were subject to acute ethanol (10% v/v) over 120-minute time course. Cell lysates (input) and purified sumoylated proteins (SUMO conjugates) were examined via Western analysis using an anti-HSV antibody to detect Smc5 and Smc6. (D) Fold change sumoylation of Smc5 and Smc6 in 10% ethanol. Each value represents the mean and SEM values are indicated as bars (*n* = 3), one-way ANOVA with Bonferroni *post hoc* test was used to compare mutants vs. control, significant differences (*P*<0.05) are indicated (**).

Proteins were classified as being sumoylated in response to ethanol stress if the summed peptide counts from the 5-minute and 60-minute replicates exceeded the 0-minute peptide counts by 3-fold or greater and were statistically significant (p>0.050) in their changes ((Oeser et al., 2016) and Supplemental Table 1)). After the MS analysis, we found 18 proteins that appear to be significantly sumoylated during acute ethanol exposure (Table 1), with 83.33% of the proteins (15 out of 18) identified as chromatin-binding and 38.89% (7 out of 18) of proteins identified as chromatin-binding transcription factors by a Gene Ontology (GO) analysis (Table 2: Gcr1, Tec1, Hap1, Ste12, Cst6, Met4, and Upc2). Of the 18 proteins identified, the top three proteins were the chromatin structural proteins Top2, Smc5, and Smc6 (Fig 2B). We were most intrigued by Smc5 and Smc6 because they form a highly conserved complex with a known role in DNA replication and repair (Duan et al., 2009b; Gallego-Paez et al., 2014; Gill, 2004; Irmisch et al., 2009; Menolfi et al., 2015; Tsuyama et al., 2006).

**Table 1.**
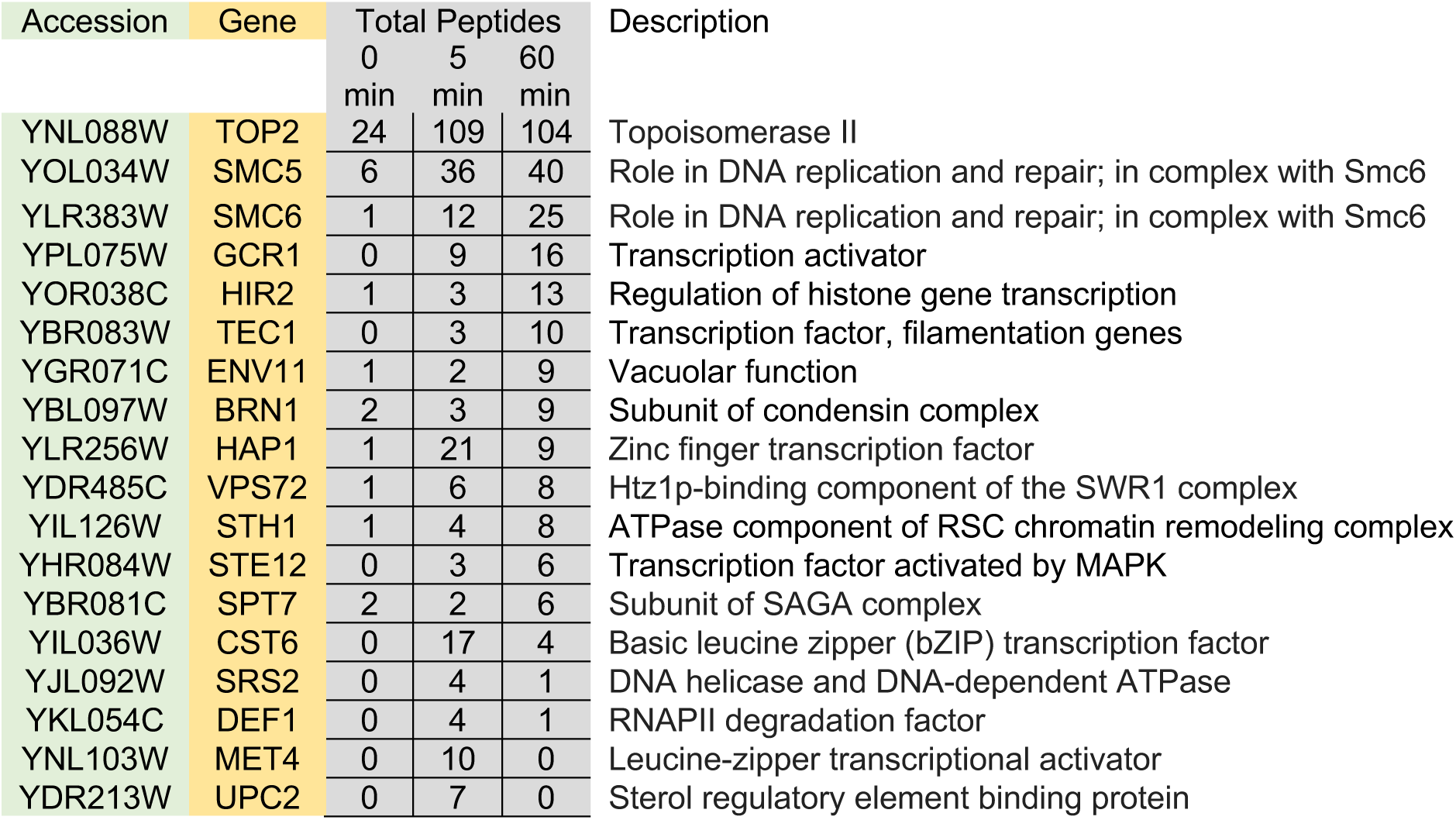
Proteins sumoylated during acute ethanol exposure

**Table 2.** Gene Ontology Analysis

| GO Terms from the molecular function Ontology |  |  |  |
| --- | --- | --- | --- |
| GO Term (GO ID) | Genes Annotated | GO Term Usage | Genome Frequency |
| DNA binding (GO:0003677) | YBR083W, YDR213W, YGR071C, YHR084W, YIL036W, YIL126W, YJL092W, YKL054C, YLR256W, YLR383W, YNL027W, YNL088W, YOL034W, YOR038C, YPL075W | 15 of 18 genes, 83.33% | 602 of 6411 annotated genes, 9.39% |
| DNA-binding transcription factor activity (GO:0003700) | YBR083W, YDR213W, YHR084W, YIL036W, YLR256W, YNL103W, YPL075W | 7 of 18 genes, 38.89% | 180 of 6411 annotated genes, 2.81% |

We confirmed Smc5/6 were sumoylated during ethanol stress by enriching for sumoylated proteins from His_6_-FLAG-*SMT3* cell lysates. To do this, we created 3xHSV tagged versions of Smc5 and Smc6 that were expressed from their endogenous promoters. By Western analysis, each protein demonstrated multiple higher molecular weight SUMO conjugates after 10% (v/v) ethanol stress over a 2-hour period (Fig 2C). We chose to verify Smc5/6 sumoylation over a 2-hour time course to confirm that sumoylation remained stable in 10% ethanol. We next quantified sumoylation of Smc5 and Smc6 over the same 2-hour period and found that Smc5 had a greater fold change increase in sumoylation over time compared to Smc6 (Fig 2D). Even though we confirmed that ethanol stress induced sumoylation of our top 3 hits (Smc5 and Smc6; Top2, data not shown), we specifically chose to pursue Smc5 and Smc6 ethanol-induced sumoylation for these studies because their sumoylation during acute ethanol stress has not yet been reported.

As an additional analysis, we expressed 3xHSV-Smc5 in a yeast strain in which all lysine residues of the *SMT3* gene itself have been mutated to arginine residues (Esteras et al., 2017), thus allowing us to determine if the ethanol-induced sumoylation of Smc5 is due to multiple mono-sumoylation events or is a chain of poly-sumoylation. In the case of Smc5, there was a predominantly single sumoylation pattern when expressed in the mutated *SMT3* strain (Supplemental Fig 1). However, we did observe slightly higher sumoylated species that indicates there might be additional mono-sumoylation sites if SUMO chain formation is eliminated. While identification of all Smc5 and 6 sumoylation sites is of interest for future studies, here we chose to examine the broader features of ethanol-induced Smc5/6 sumoylation.

### The E3 SUMO ligase Mms21 promotes the sumoylation of Smc5/6 during acute ethanol stress

After identifying Smc5 and Smc6 as sumoylation targets during acute ethanol stress, we wanted to determine the enzymes required for their ethanol-induced sumoylation. Unlike the ubiquitination system, which has over 100 E3 ubiquitin ligases in yeast (De Bie and Ciechanover, 2011), the sumoylation system only has four known E3 SUMO ligases: Siz1, Siz2, Cst9, and Mms21 (Gill, 2004; Hay, 2001). It has been previously reported that the E3 for Smc5/6 is Mms21 during DNA damage (Bermúdez-López et al., 2015; Duan et al., 2009a; Liang et al., 2018), but it was unclear if this was the case during acute ethanol exposure. *SIZ1, SIZ2*, and *CST9* are not essential genes and can be deleted. *MMS21* is essential and we therefore used an auxin-inducible degron (AID) depletion strategy (Havens et al., 2012; Nishimura et al., 2009), wherein Mms21 was fused to an auxin degron and its depletion is induced by addition of auxin to the media. We found that, after Mms21 depletion by addition of auxin for 90 minutes, Smc5/6 sumoylation decreased approximately 75% during the 60-minute time course of acute ethanol stress (Fig 3A), consistent with the similar reduction in Mms21 levels. We also investigated if the non-essential E3s were involved in sumoylation induced by acute ethanol exposure. We found that complete loss of Siz1, Siz2, or Cst9 did not significantly reduce Smc5/6 sumoylation during acute ethanol stress (Fig 3B).

**Figure 3.**
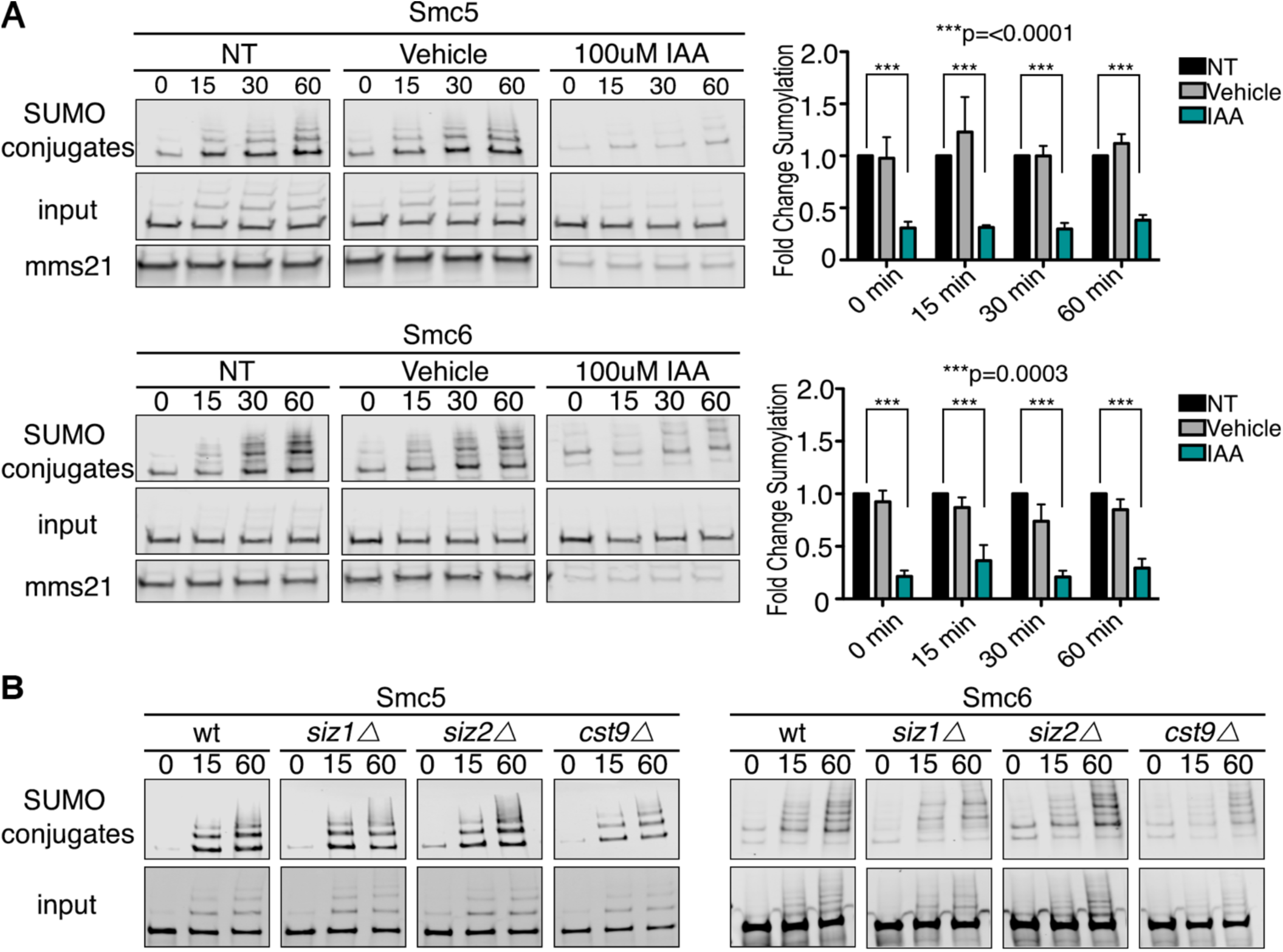
The E3 SUMO ligase Mms21 promotes sumoylation of Smc5/6 during ethanol stress. (A) Cells expressing His_6_-FLAG-*SMT3, MMS21*-3HSV-AID and 3xHSV tagged Smc5 or Smc6 from their endogenous promoters were grown to early log-phase then treated with NT, Vehicle, or auxin (IAA) for 90 minutes before exposure to acute ethanol for 60 minutes. Cell lysates (input) and purified sumoylated proteins (SUMO conjugates) were examined via Western analysis with an anti-HSV antibody to detect Mms21, Smc5 and Smc6. Fold changes in sumoylation were calculated utilizing values from Image Studio Lite software. Values were normalized to inputs and NT samples set to a value of 1.0. Error bars show SEM for triplicate samples. NT samples compared to IAA were significantly different in Smc5 and Smc6 with ***p=<0.0001 and ***p=0.0003 respectively. (B) Wild-type (WT), *siz1Δ, siz2Δ*, or *cst9Δ* cells expressing His_6_-FLAG-SMT3 and either 3xHSV Smc5 or Smc6 were subjected to 60-minutes of acute ethanol (10% v/v) exposure. Cell lysates (input) and metal affinity purified sumoylated proteins (SUMO conjugates) were examined by Western analysis with an anti-HSV antibody.

### Ethanol exposure leads to Smc5/6 sumoylation in G1 and G2 phases that is reduced in S phase

Sumoylation of proteins is essential for progression through the cell cycle and SUMO conjugation to its targets dynamically change during different stages under normal conditions (Talamillo et al., 2020). A common feature of yeast cells is that exposure to stress often affects progression through the cell cycle (Jorgensen and Tyers, 2004). For example, yeast cells have been shown to halt in the G1 phase of the cell cycle after exposure to stress (Qu et al., 2019). This pause is generally thought to allow cells the time to resolve or adapt to the stress before proceeding through the cell cycle (Qu et al., 2019).

Chromatin undergoes regulated changes during the cell cycle (Antonin and Neumann, 2016; Ma et al., 2015). Since the Smc5/6 complex is chromatin-associated and its chromatin localization is altered specifically during S phase (Jeppsson et al., 2014), we were interested in determining the stages of the cell cycle where Smc5/6 undergoes ethanol-dependent sumoylation. We first examined if Smc5/6 could be sumoylated during G1-phase arrest induced by addition of alpha factor. We found that, when compared to controls, arrest in G1 allowed the same sumoylation as unarrested cells after acute ethanol exposure (Fig 4B). We next examined Smc5/6 sumoylation in cells arrested in G2 phase by addition of nocodazole (ND). Again, we found that G2 arrest also allowed Smc5/6 sumoylation similar to unarrested cells (Fig 4C). Finally, we arrested cells in S phase using hydroxyurea (HU). In this case, ethanol-induced Smc5/6 sumoylation was considerably reduced (Fig 4D). To determine that HU addition did not block global sumoylation in the cell, we also examined sumoylation of the transcriptional co-repressor Cyc8 during HU treatment and hyperosmotic stress (1.2M sorbitol), which is a condition we previously found Cyc8 to be sumoylated (Oeser et al., 2016). We found that hyperosmotic-stress induced Cyc8 sumoylation still occurred after S-phase arrest (Fig 4E), thereby ruling out an inhibitory effect of HU addition on global sumoylation. Altogether, Smc5/6 are sumoylated during alpha factor and nocodazole arrests in G1 and G2/M phases respectively but are not sumoylated during HU-arrest in S phase.

**Figure 4.**
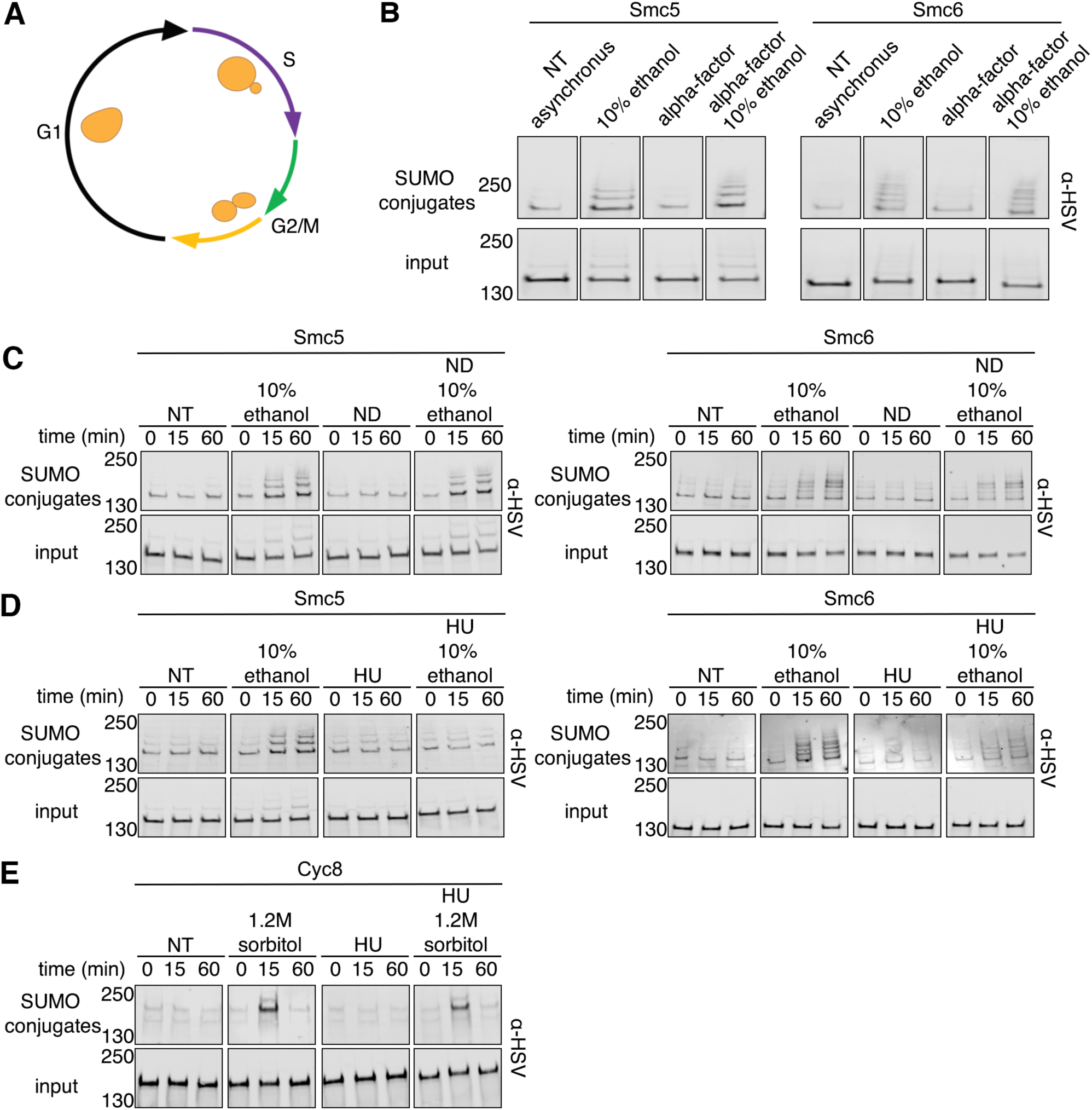
Ethanol exposure leads to Smc5/6 sumoylation in G1 and G2 phases that is reduced in S phase. (A) Schematic of cell cycle with cell morphology for each phase. (B) Asynchronous cells expressing His_6_-FLAG-*SMT3* and either 3xHSV-tagged Smc5 or Smc6 from their endogenous promoters were arrested in G1 with *α*-factor for 90-minutes prior to treatment with 10% ethanol for 60-minutes. Cell lysates (input) and purified sumoylated proteins (SUMO conjugates) were examined by Western analysis with an anti-HSV antibody. (C and D) Similar experiments to (B) where asynchronous cells were arrested in G2/M with nocodazole and S-phase with HU respectively for 90-minutes prior to exposure to 10% ethanol for a 60-minute time course. (E) Asynchronous cells expressing His_6-_FLAG-SMT3 and 3xHSV Cyc8 from its endogenous promoter were arrested in S-phase with HU for 90-minutes prior to 60-minutes of hyperosmotic (1.2M sorbitol) stress. Cell lysates (input) and purified sumoylated proteins (SUMO conjugates) were examined by Western analysis using an anti-HSV antibody.

### Smc5/6 are sumoylated during heat stress and DNA damage, but not hyperosmotic stress

We previously reported that global sumoylation patterns and kinetics differ between distinct stress conditions (Oeser et al., 2016). To gain better insight into the functional purpose of Smc5 and Smc6 sumoylation, we examined their sumoylation patterns during heat stress (42°C), hyperosmotic stress (1.2M sorbitol), and DNA damage (exposure to methyl methanesulfonate (MMS)). After exposing cells to 42°C heat shock over a time course of 60 minutes, we found that Smc5 and Smc6 SUMO conjugates accumulated at a slower rate compared to exposure to acute ethanol (Fig 5A), though the pattern of sumoylation banding was similar. We also subjected cells to hyperosmotic stress (1.2M sorbitol) over 60 minutes and found that there was a rapid decrease in sumoylation after 5 minutes before an equally rapid increase at 15 minutes that remained stable for the duration of the time course (Fig 5B). The rapid decrease in Smc5/6 sumoylation is consistent with what we previously observed during hyperosmotic stress; proteins are desumoylated to provide a free pool of SUMO that is readily available for Tup1 and Cyc8 sumoylation (Oeser et al., 2016). There is an increase in Smc5/6 sumoylation following the initial decrease during hyperosmotic stress; however, we do not think Smc5/6 are sumoylated in response to hyperosmotic stress. Instead, we think that we are likely observing the return to basal sumoylation prior to hyperosmotic stress as the cells adapt to hyperosmotic stress and Tup1/Cyc8 are desumoylated (Oeser et al., 2016).

**Figure 5.**
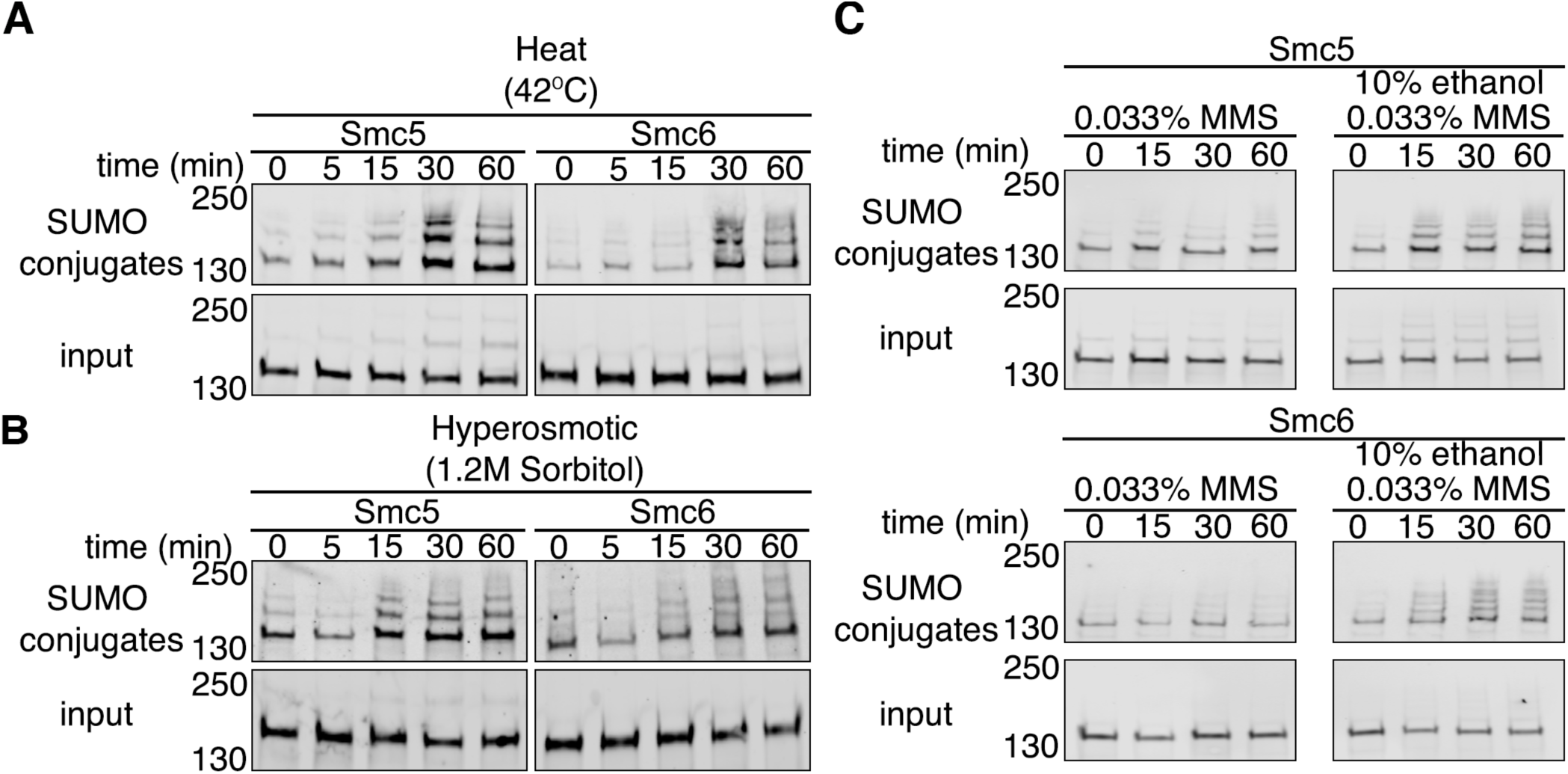
Smc5/6 are sumoylated during heat shock and MMS stress, but not hyperosmotic. (A-B) Cells expressing His_6_-FLAG-*SMT3* and 3xHSV epitope tagged Smc5 or Smc6 from their endogenous promoters were exposed to either heat (42°C) or hyperosmotic (1.2M sorbitol) stress over a 60-minute time course. Cell lysates (input) and metal affinity purified sumoylated proteins (SUMO conjugates) were examined by Western analysis using an anti-HSV antibody. (C) Similar experiment to (A) but cells were subjected to 0.033% MMS over 60 minutes with or without 10% ethanol.

It has been reported that Smc5 and Smc6 are sumoylated after exposure to levels of methyl methanesulfonate (MMS) that induce DNA damage (Zapatka et al., 2019). We wanted to verify that Smc5 and Smc6 were sumoylated during a time course of MMS exposure, and if the addition of ethanol with MMS treatment caused no effect or was additive. We predicted that if both had the same effect, that addition of ethanol would not change Smc5 or Smc6 sumoylation beyond MMS treatment. We exposed cells to 0.033% MMS and found that the sumoylation of Smc5 and Smc6 increased over time with acute MMS exposure (Fig 5C, left panels). However, addition of ethanol with MMS caused significantly more Smc5/6 sumoylation than MMS treatment alone (Fig 5C, right panels). Thus, it appears that MMS and ethanol might have different physiological effects that lead to additive Smc5/6 sumoylation.

### Acute ethanol stress causes DNA damage

Treatment of yeast cells with MMS causes DNA damage through alkylation of guanine and adenine moieties (Beranek, 1990). Chronic, long-term treatment of yeast cells with ethanol has recently been reported to increase DNA mutation rates through error-prone DNA polymerases being recruited to the replication fork (Voordeckers et al., 2020). Thus, we wanted to explore if acute ethanol exposure led to DNA damage. *In vivo* ways to examine DNA damage have been developed through observing foci formed by the homologous recombination protein Rad52 fused to tdTomato (Estrem and Moore, 2019), and phosphorylation of the intra-S phase checkpoint kinase Rad53 (Sanchez et al., 1996). We examined cells that were either untreated, treated with 0.033% MMS or 10% ethanol alone, or 0.033% MMS plus 10% ethanol together. Compared to no treatment, 15 minute treatment of cells with either 10% ethanol or 0.033% MMS caused the formation of Rad52 foci in about 1% of cells in an asynchronous cell culture (Figure 6A-B), and this is consistent with that observed in prior studies examining MMS treatment (Estrem and Moore, 2019). There was no increase in Rad52 foci frequency in cells treated with both 10% ethanol and 0.033% MMS (Figure 6A-B). By this measure, ethanol and MMS have a similar effect with inducing Rad52 foci. Interestingly, acute ethanol treatment did not induce phosphorylation of Rad53 whereas MMS did cause Rad53 phosphorylation (Fig 6C), indicating that acute ethanol exposure does not trigger the intra-S phase checkpoint like MMS exposure (Barlow and Rothstein, 2010). Importantly, addition of ethanol and MMS together did not trigger Rad53 phosphorylation, suggesting that ethanol exposure causes effects that are epistatic to MMS exposure. Altogether, we conclude that acute ethanol exposure does lead to DNA damage, but it did not appear to additionally increase DNA damage when cells were treated together with ethanol and MMS.

**Figure 6.**
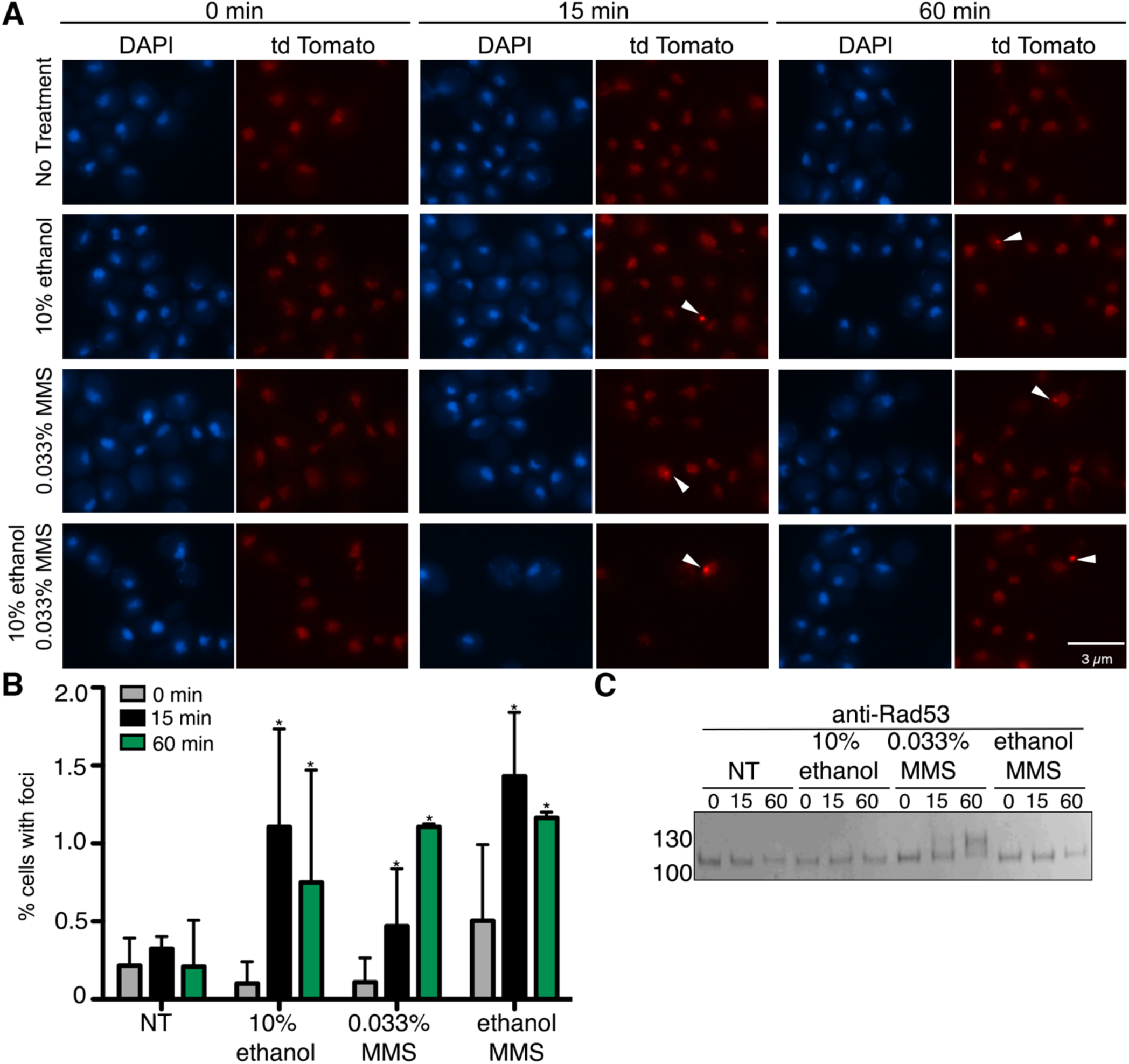
Ethanol causes formation of Rad52 foci but not Rad53 phosphorylation. (A) Rad52-tdTomato cells were exposed to acute ethanol (10%), 0.033% MMS, or combined ethanol and MMS over a 60-minute time course. Cells fixed at indicated timepoints and imaged by fluorescent microscopy. Five fields of cells for each condition with ≥40 cells/field were counted for the presence of nuclear foci (B). (C) Cells were exposed to acute ethanol (10%), 0.033% MMS, or combined ethanol and MMS over a 60-minute time course. Rad53 phosphorylation was observed by Western analysis using and anti-Rad53 antibody.

## Discussion

Adaptation to stress is essential for cellular survival, and the cell utilizes distinct multifaceted approaches to re-establish homeostasis during particular stress conditions. Without undergoing stress adaptation, prolonged cellular stress can lead to the irreversible damage of cellular components that can ultimately impact cell viability (Figure 7). In the case of ethanol, chronic exposure to high concentrations of ethanol leads to alterations in membrane fluidity and lipid composition, increased production of reactive oxygen species (ROS) through altering oxidative phosphorylation in the mitochondria, and cause protein denaturing and misfolding (Auesukaree, 2017; Kato et al., 2011, 2019). In this study, we chose to examine proteins that become sumoylated during acute ethanol exposure. Of the 18 proteins we identified in a mass spectrometric analysis that become increasingly sumoylated during acute ethanol stress, we found that 15 of the 18 proteins are known to be chromatin-associated proteins (Table 2). This is not unexpected as protein sumoylation is known to regulate multiple chromatin-associated processes including the DNA damage checkpoint (Munk et al., 2017; Wu et al., 2014), regulation of chromosome structure (Nacerddine et al., 2005; Tanaka et al., 1999), and chromosome movement (Seeber and Gasser, 2017; Zhao, 2018).

**Figure 7.**
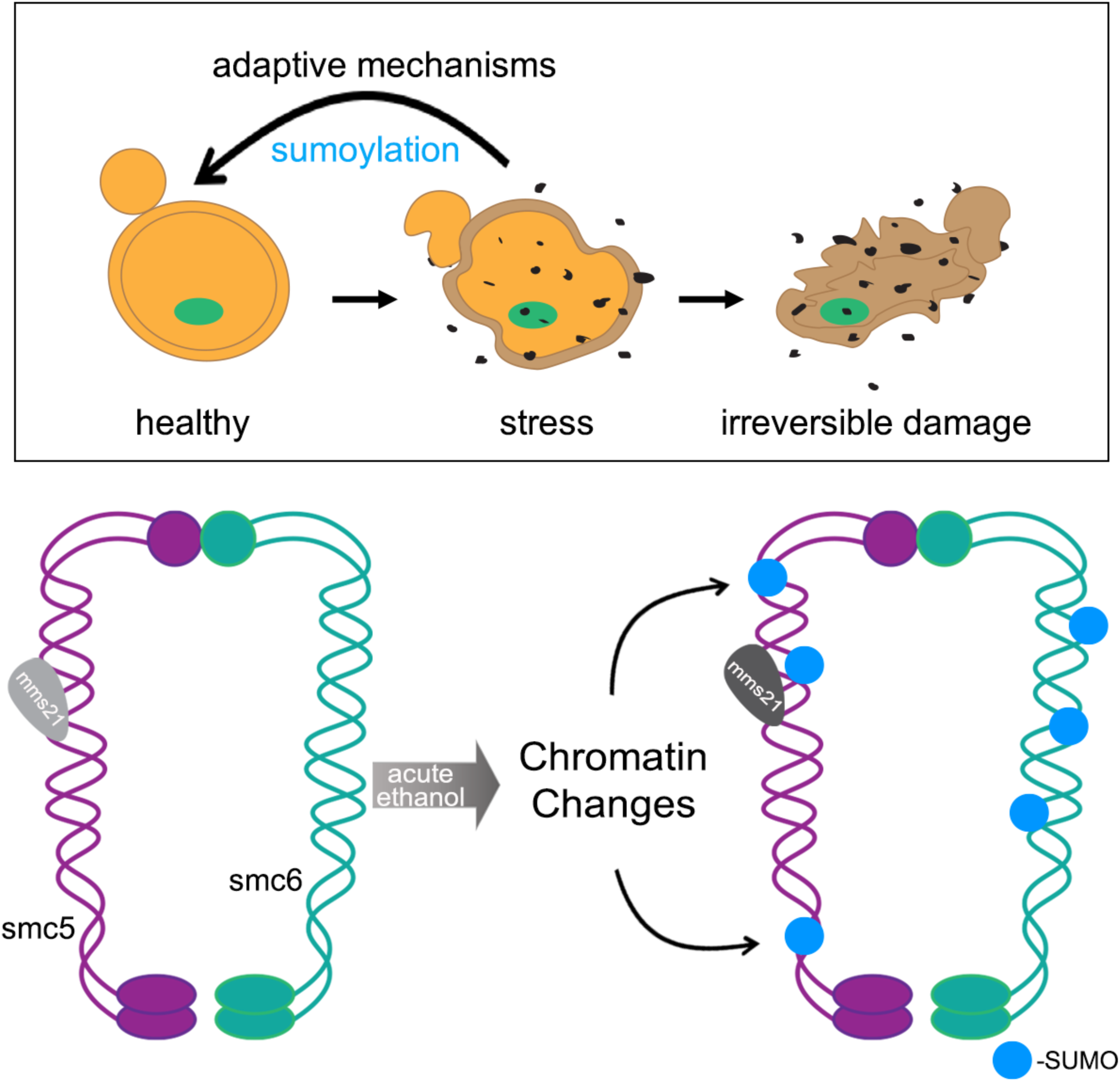
Model for cellular stress and Smc5/6 sumoylation.

Although not the main focus of this study, we think it is important to comment on the scope of proteins identified in the MS analysis. Of the 18 proteins discovered, 7 are known to function as transcription factors (Table 1 and 2: Gcr1, Tec1, Hap1, Ste12, Cst6, Met4, and Upc2). The particular functions of these transcription factors reflect the ways in which ethanol alters cellular physiology. The cellular effects of ethanol exposure include alterations in membrane fluidity and lipid composition (Tóth et al., 2014), changes in glucose and amino acid uptake (Yang et al., 2012), a reduction in the activities of glycolytic enzymes (Tóth et al., 2014), and disruption of membrane integrity (Stanley et al., 2010). In terms of membrane fluidity, Upc2 is a transcription factor that undergoes regulated cleavage from the ER membrane to activate sterol biosynthesis genes (Joshua and Höfken, 2017), and it is known that ethanol exposure leads to the increased synthesis and presence of unsaturated fatty acids and ergosterol in the membrane (Henderson and Block, 2014). For changes in glucose and amino acid uptake, Gcr1 regulates genes involved in glycolysis (Hossain et al., 2016), and Met4 regulates genes in the sulfur amino acid biosynthesis pathway (McIsaac et al., 2012). Tec1 and Ste12 are involved in regulating filamentous and invasive growth pathways (Mayhew and Mitra, 2014). Cst6 is also known to regulate stress and carbon utilization pathways (Pohlers et al., 2017), and deletions are sensitive to ethanol stress (Liu et al., 2016). We have confirmed Cst6 is sumoylated under acute ethanol stress (unpublished data). The remaining transcription factors have yet to be verified for ethanol-induced sumoylation by our group, but Tec1, Hap1, Met4, and Upc2 have been identified in a general screen for sumoylated proteins (Esteras et al., 2017). Altogether, it is notable that the cellular processes affected by ethanol exposure are regulated by transcription factors that are sumoylated during acute ethanol stress. Alteration of transcription factor activity in these pathways is consistent with yeast cells mounting an adaptative response to manage the cellular dysfunction that occurs with exposure to ethanol. Further studies are needed to understand the transcriptional responses that might occur through ethanol-induced sumoylation of transcription factors and how sumoylation alters their function.

In addition to transcription factors, major proteins that showed increased sumoylation upon ethanol exposure were Smc5, Smc6, and Top2, which are known to form a highly conserved chromatin structure complex (Aragón, 2018). The Smc5/6 complex is one of four highly conserved <u>s</u>tructural <u>m</u>aintenance of <u>c</u>hromosomes (SMC) complexes found in eukaryotes and is best known for its role in DNA repair and overall genome stability (Aragón, 2018). It has been described that Smc5/6 sumoylation occurs as a regulatory consequence of collapsed replication forks and has a functional role in modulating replication associated repair and error free DNA bypass via the Mph1 helicase (Zapatka et al., 2019). The Smc5/6 complex also has been shown to interact with the E3 SUMO ligase Mms21 to promote the sumoylation of the STR helicase complex that acts in the removal of recombination intermediates (Bonner et al., 2016); however we did not find members of the complex Sgs1, Top3, or Rmi1 to undergo increased sumoylation in our MS analysis (Supplemental Table 1). We did find that Smc5/6 are sumoylated during arrest of cells in the G1 and G2/M phases of the cell cycle after acute ethanol exposure (Fig 4), but we did not observe Smc5/6 sumoylation during S phase arrest induced by addition of HU. We think this indicates that there are specific windows when ethanol-induced Smc5/6 sumoylation occurs. Considering that Smc5/6 have nearly identical chromatin localizations during G1 and G2/M phases but different ones during S phase (Jeppsson et al., 2014), it may be that chromatin context is important for Smc5/6 sumoylation during ethanol stress.

We also observed an increase in Smc5/6 sumoylation after exposure of cells to both ethanol and the DNA alkylating agent MMS (Fig 5C), indicating that acute ethanol stress induced Smc5/6 sumoylation may operate through a different mechanism than MMS stress alone. We found that both ethanol and MMS exposure induced Rad52 foci to similar extent (Fig 6A-B), leading us to conclude that both might lead to Smc5/6 sumoylation through a DNA damage response, though the nature of the damage might be different. In fact, ethanol exposure did not trigger the intra-S phase checkpoint as seen by the phosphorylation of Rad53 whereas MMS treatment did lead to Rad53 phosphorylation (Figure 6C). Currently, the function of ethanol-induced Smc5/6 sumoylation is not clear. Future experiments will be needed to determine if Smc5/6 sumoylation is due to increased DNA damage, altered chromatin structure, or a response to protein misfolding that could lead to both DNA damage and/or chromatin structural loss. Consistent with this idea, we note that heat shock also induced Smc5/6 sumoylation, and heat shock can also lead to similar changes in protein/chromatin structure as well as DNA damage (Niskanen and Palvimo, 2017).

Overall, we conclude from the data presented here that there are two responses the cell elicits during acute ethanol exposure through sumoylation: one is to protect chromatin structure and the other is to mount an adaptive response through altered gene transcription. We previously found that sumoylation modulates a transient phase separation in the Tup1-Cyc8 transcriptional co-repressor complex (Oeser et al., 2016), indicating a chromatin-modifying activity for sumoylation during hyperosmotic stress. From a transcriptional perspective, we know from our previous work (Nadel et al., 2019; Oeser et al., 2016) and other studies (Stielow et al., 2008; Zhao, 2018; Zhou et al., 2004) that genes involved either directly in transcription or its modulation have been reported to be sumoylated. How stress-induced sumoylation affects protein activity, localization, or stability remains an open question in the field, and it may be dictated by the magnitude and duration of the stress. Our studies indicate that transient sumoylation targets chromatin-associated proteins during stress adaptation, and support that idea that transient sumoylation is a common regulatory phenomenon during stress conditions.

## Acknowledgements

We thank Jeffrey Moore (University of Colorado Anschutz Medical Campus) for the Rad52-tdTomato strain. We thank Luis Aragón (MRC Clinical Sciences Centre, Imperial College, London) for the yeast strain CCG9474 that allowed for identification of SUMO chains. We thank Matt Kaeberlein’s lab (University of Washington) for experimental assistance with the growth curves. We thank Xiaolan Zhao (Sloan Kettering Institute) and Sue Biggins (Fred Hutchinson Cancer Research Center) for critical advice. This work was supported by a NIH/NGIMS training grant T32GM007270 (A.I.B) a HHMI Gilliam fellowship GT10825 (A.I.B), a NIH/NIGMS grant R01AG031136 (R.G.G.), NIH/NIGMS grants R01GM114112 and R35GM136234 (R.G.G.). The authors declare no competing financial interests.

## Materials and Methods

### Yeast strains and plasmids

Yeast strains and plasmids used in this study are listed in (Table 3). Standard yeast genetic methods were used for this study (Guthrie C, 1991). All gene deletions were verified by colony PCR and phenotypic analyses when available.

**Table 3:** yeast strains.

| **Strain** | **Genotype** | **Reference** |
| --- | --- | --- |
| RGY 5266 | *met15Δ0, his3Δ1, ura3Δ0, leu2Δ0, 6His-FLAG-SMT3::HIS3MX* | Oeser et al., 2016 |
| RGY 6005 | *met15Δ0, his3Δ1, ura3Δ0, leu2Δ0, 6His-FLAG-SMT3::HIS3MX, Smc6-3HSV::KanMX* | this study |
| RGY 6014 | *met15Δ0, his3Δ1, ura3Δ0, leu2Δ0, 6His-FLAG-SMT3::HIS3MX, Smc5-3HSV::KanMX* | this study |
| RGY 6339 | *met15Δ0, his3Δ1, ura3Δ0, leu2Δ0, 6His-FLAG-SMT3::HIS3MX, MMS21-3HSV-IAA1.T10::KanMX, AFB2::LEU2, Smc5-3HSV::URA3* | this study |
| RGY 6340 | *met15Δ0, his3Δ1, ura3Δ0, leu2Δ0, 6His-FLAG-SMT3::HIS3MX, MMS21-3HSV-IAA1.T10::KanMX, AFB2::LEU2, Smc6-3HSV::URA3* | this study |
| RGY 6346 | *met15Δ0, his3Δ1, ura3Δ0, leu2Δ0, 6His-FLAG-SMT3::HIS3MX, siz2Δ::KanMX, Smc5-3HSV::URA3* | this study |
| RGY 6347 | *met15Δ0, his3Δ1, ura3Δ0, leu2Δ0, 6His-FLAG-SMT3::HIS3MX, cst9Δ::KanMX, Smc6-3HSV::URA3* | this study |
| RGY 6361 | *met15Δ0, his3Δ1, ura3Δ0, leu2Δ0, 6His-FLAG-SMT3::HIS3MX, siz1Δ::KanMX, Smc5-3HSV::URA3* | this study |
| RGY 6362 | *met15Δ0, his3Δ1, ura3Δ0, leu2Δ0, 6His-FLAG-SMT3::HIS3MX, siz1Δ::KanMX, Smc6-3HSV::URA3* | this study |
| RGY 6363 | *met15Δ0, his3Δ1, ura3Δ0, leu2Δ0, 6His-FLAG-SMT3::HIS3MX, siz2Δ::KanMX, Smc6-3HSV::URA3* | this study |
| RGY 6344 | *met15Δ0, his3Δ1, ura3Δ0, leu2Δ0, 6His-FLAG-SMT3::HIS3MX, cst9Δ::KanMX, Smc5-3HSV::URA3* | this study |
| yJM2468 | *ura3-52, lys2-801, leu2-Δ1, his3-Δ200, trp1-Δ63, RAD52-tdTomato::HIS3MX* | Estrem and Moore, 2019 |

### Growth and stress conditions

Cells were grown to a density of ∼1.5×10^7^ cells/ml at 30°C in yeast extract peptone dextrose (YPD) media prior to stress induction. All 0 time point samples were collected before stress induction. For ethanol stress, volume per volume amounts were added to cultures for a final concentration of 10% v/v ethanol. For hyperosmotic stress, equal volumes of culture and YPD+2.4M sorbitol were combined for a final concentration of 1.2M sorbitol. For heat stress, cells were pelleted and resuspended in YPD media warmed to 42°C and placed in a shaking platform 42°C incubator.

### Sumoylated protein purification

50ml aliquots of cells were collected at designated time points after stress and flash frozen in liquid nitrogen. Harvested cells were lysed by vortexing with acid washed glass beads at 4°C in 1ml denaturing lysis buffer (8M urea, 50mM Tris pH 8.0, 0.05% SDS with 2mM PMSF and 20mM NEM). Aliquots representing 5% of the input were set aside. Cell lysates were incubated with TALON resin (Novagen) overnight at 4°C. Resin was washed 3x with wash buffer (8M urea, 50mM Tris pH 8.0, 200mM NaCl, 0.05% SDS). Sumoylated proteins were eluted from beads with loading buffer (8M urea, 10mM MOPS, 10mM EDTA, 1% SDS, 0.01% bromophenol blue, pH 6.8) and incubated at 65°C for 10 minutes.

### Western analysis

Total cell lysates and purified sumoylated proteins were resolved by SDS-PAGE using 4-20% gradient gels (BioRad). Western analyses were performed with mouse anti-FLAG (1:2500, Sigma), mouse anti-HSV (1:2500, Novagen), rabbit anti-HSV (1:2000, Abcam), and rabbit anti-Rad53 (1:2000, Abcam).

### Mass spectrometry analyses

Sumoylated proteins from cells exposed to 0, 5, or 60 minutes of 10% (v/v) ethanol stress were enriched by metal affinity chromatography as described above. Samples were run 1 cm into a 4-20% SDS-PAGE gel and bands were excised. Proteins in gel slices were digested with trypsin and digestion products desalted and dried by vacuum centrifugation. Dried peptide mixtures were resuspended in 7ul of 0.1% formic acid. 5μL was analyzed using a LTQ OrbiTrap mass spectrometer (Thermo Scientific). Complete MS methods were performed as previously described (Richardson et al., 2012).

The protein database search algorithm X!Tandem (Craig and Beavis, 2004) was used to identify peptides from the Saccharomyces Genome Database (http://www.yeastgenome.org). Peptide false discovery rates were measured using Peptide Prophet (Keller et al., 2002). Identified peptides were filtered using Peptide Prophet scores of ≥0.55 (∼10% false discover rate). The entire dataset is in Table S1). The significance of the changes in peptide counts between 0, 5, 60 minutes of ethanol stress was determined by a two-tailed, homoscedastic student’s t-test. Data was filtered by a p≤0.01 and a +/-3-fold change in summed peptide counts. Final filtered data is in Table 1.

### Cell cycle analysis

Asynchronous cells were diluted back to a density of ∼0.385×10^7^ cells/ml in YPD or YPD+1% DMSO (nocodazole) media prior to cell cycle halt and stress induction. For G1 arrest, 50μg/ml alpha-factor (Sigma) was added immediately to cultures and incubated at 30°C for 90 minutes. For G2/M arrest, cells were incubated at 30°C for 2 hours then treated with 0.05 mg/ml nocodazole (Sigma) for an additional hour. For S arrest, 100mM hydroxyurea (Sigma) was added immediately to cultures and incubated at 30°C for 90 minutes. All cell arrests were verified by pelleting a 200ul aliquot of culture and examining under a phase contrast microscope (Nikon). ∼75-90% of cells per field were observed to be arrested at a given phase. For G1 arrest cells were observed to be either large and unbudded or with shmoo morphology. Cells arrested in S-phase were observed to be large with small buds. While G2/M arrested cells were observed as dumbbells.

### Auxin-degron depletion experiments

Cells were grown to a density of ∼0.86×10^7^ cells/ml at 30°C in Yeast Complete (YC) media prior to addition of either NT, vehicle (1:1000, 95% EtOH), or 100μM 3-Indoleacetic acid (IAA, Sigma). Cells were then incubated for 90 minutes at 30°C and prior to stress induction. 10% (v/v) ethanol was then added to cultures for an additional hour with 50ml aliquots of cells collected at designated timepoints flash frozen in liquid nitrogen before proceeding with sumoylated protein purification.

### Fluorescence microscopy

Aliquots of cells at each time point after ethanol stress were removed, fixed in 4% paraformaldehyde solution for 15 minutes at room temperature then washed with 1X PBS. Cells were imaged on a Nikon Eclipse 90i with a 100x objective, filters for GFP [HC HiSN Zero Shift filter set with excitation wavelength (450–490 nm), a dichroic mirror (495 nm) and emission filter (500–550 nm)], tdTomato [HC HiSN Zero Shift filter set with excitation wavelength (530–560 nm), a dichroic mirror (570 nm) and emission filter (590–650 nm)] or DAPI [HC HiSN Zero Shift filter set with excitation wavelength (325–375 nm), a dichroic mirror (400 nm) and emission filter (435–485 nm)], and a Photometrics Cool Snap HQ2 cooled CCD camera with NIS-Elements acquisition software.

### Image processing

All blots were scanned using a Licor Odyssey CLx and ImageStudio Lite. All images were processed with a MacBook Pro or iMac computer (Apple) using Photoshop (Adobe).

### Rigor and reproducibility

All biochemical and microbiological assays were performed in triplicate. For fluorescence microscopy, three separate researchers quantified loci formation. Statistics used were, paired students t-tests (Figure 3 and 6) and one-way ANOVA with Bonferroni *post hoc* test (Figure 2).

---

**Supplemental Figure 1.**
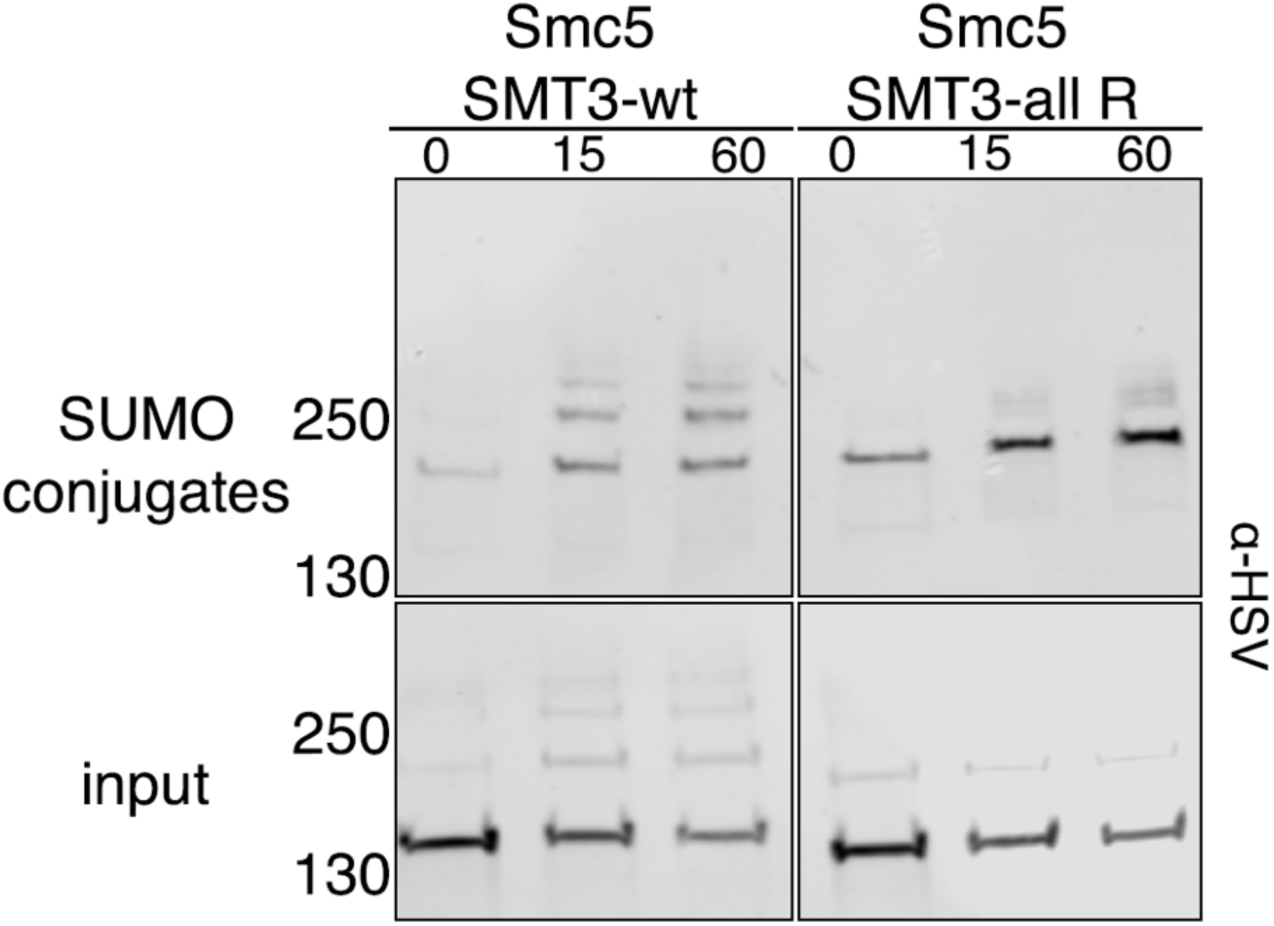
Ethanol-induced Smc5 appears to be primarily a poly-sumoylation chain. (A-B) Cells with 3xHSV-Smc5 expressed from its endogenous promoter and expressing either His_6_-*SMT3* that contains all lysine residues (SMT3-wt) or all lysine residues converted to arginine residues (SMT3-all R) were exposed to 10% ethanol over a 60-minute time course. Cell lysates (bottom panels) and metal affinity purified sumoylated proteins (top panels) were examined by Western analysis using an anti-HSV antibody.

## Notes

### Competing Interest Statement

The authors have declared no competing interest.

## References

Antonin W, Neumann H. 2016. Chromosome condensation and decondensation during mitosis. Curr Opin Cell Biol 40:15–22. doi:10.1016/j.ceb.2016.01.013

Aragón L. 2018. The Smc5/6 Complex: New and Old Functions of the Enigmatic Long-Distance Relative. Annu Rev Genet 52. doi:10.1146/annurev-genet-120417-031353

Auesukaree C. 2017. Molecular mechanisms of the yeast adaptive response and tolerance to stresses encountered during ethanol fermentation. J Biosci Bioeng 124:133–142. doi:10.1016/j.jbiosc.2017.03.009

Barlow JH, Rothstein R. 2010. Timing is everything: cell cycle control of Rad52. Cell Div 5:29– 31. doi:10.1186/1747-1028-5-7

Beranek DT. 1990. Distribution of methyl and ethyl adducts following alkylation with monofunctional alkylating agents. Mutat Res - Fundam Mol Mech Mutagen 231:11–30. doi:10.1016/0027-5107(90)90173-2

Bermúdez-López M, Pociño-Merino I, Sánchez H, Bueno A, Guasch C, Almedawar S, Bru-Virgili S, Garí E, Wyman C, Reverter D, Colomina N, Torres-Rosell J. 2015. ATPase-Dependent Control of the Mms21 SUMO Ligase during DNA Repair. PLoS Biol 13. doi:10.1371/journal.pbio.1002089

Bonner JN, Choi K, Xue X, Torres NP, Szakal B, Wei L, Wan B, Arter M, Matos J, Sung P, Brown GW, Branzei D, Zhao X. 2016. Smc5/6 Mediated Sumoylation of the Sgs1-Top3-Rmi1 Complex Promotes Removal of Recombination Intermediates. Cell Rep. doi:10.1016/j.celrep.2016.06.015

Craig R, Beavis RC. 2004. TANDEM: Matching proteins with tandem mass spectra. Bioinformatics 20:1466–1467. doi:10.1093/bioinformatics/bth092

De Bie P, Ciechanover A. 2011. Ubiquitination of E3 ligases: Self-regulation of the ubiquitin system via proteolytic and non-proteolytic mechanisms. Cell Death Differ 18:1393–1402. doi:10.1038/cdd.2011.16

Duan X, Sarangi P, Liu X, Rangi GK, Zhao X, Ye H. 2009a. Structural and Functional Insights into the Roles of the Mms21 Subunit of the Smc5/6 Complex. Mol Cell 35:657–668. doi:10.1016/j.molcel.2009.06.032

Duan X, Yang Y, Chen YH, Arenz J, Rangi GK, Zhao X, Ye H. 2009b. Architecture of the Smc5/6 complex of Saccharomyces cerevisiae reveals a unique interaction between the Nse5-6 subcomplex and the hinge regions of Smc5 and Smc6. J Biol Chem. doi:10.1074/jbc.M809139200

Esteras M, Liu IC, Snijders AP, Jarmuz A, Aragon L. 2017. Identification of sumo conjugation sites in the budding yeast proteome. Microb Cell 4:331–341. doi:10.15698/mic2017.10.593

Estrem C, Moore JK. 2019. Astral microtubule forces alter nuclear organization and inhibit DNA repair in budding yeast. Mol Biol Cell 30:2000–2013. doi:10.1091/mbc.E18-12-0808

Fulda S, Gorman AM, Hori O, Samali A. 2010. Cellular stress responses: Cell survival and cell death. Int J Cell Biol. doi:10.1155/2010/214074

Gallego-Paez LM, Tanaka H, Bando M, Takahashi M, Nozaki N, Nakato R, Shirahige K, Hirota T. 2014. Smc5/6-mediated regulation of replication progression contributes to chromosome assembly during mitosis in human cells. Mol Biol Cell. doi:10.1091/mbc.E13-01-0020

Galluzzi L, Yamazaki T, Kroemer G. 2018. Linking cellular stress responses to systemic homeostasis. Nat Rev Mol Cell Biol. doi:10.1038/s41580-018-0068-0

Geiss-Friedlander R, Melchior F. 2007. Concepts in sumoylation: A decade on. Nat Rev Mol Cell Biol. doi:10.1038/nrm2293

Gill G. 2004. SUMO and ubiquitin in the nucleus: different functions, similar mechanisms? Genes Dev 18:2046–2059. doi:10.1101/gad.1214604

Guo C, Henley JM. 2014. Wrestling with stress: Roles of protein SUMOylation and deSUMOylation in cell stress response. IUBMB Life. doi:10.1002/iub.1244

Guthrie C FG. 1991. Guide to yeast genetics and molecular biology. Methods in Enzymology. doi:10.1016/0962-8924(92)90020-n

Havens KA, Guseman JM, Jang SS, Pierre-Jerome E, Bolten N, Klavins E, Nemhauser JL. 2012. A synthetic approach reveals extensive tunability of auxin signaling. Plant Physiol 160:135–142. doi:10.1104/pp.112.202184

Hay RT. 2001. Protein modification by SUMO. Trends Biochem Sci 26:332–333. doi:10.1016/S0968-0004(01)01849-7

Henderson CM, Block DE. 2014. Examining the role of membrane lipid composition in determining the ethanol tolerance of saccharomyces cerevisiae. Appl Environ Microbiol 80:2966–2972. doi:10.1128/AEM.04151-13

Hossain MA, Claggett JM, Edwards SR, Shi A, Pennebaker SL, Cheng MY, Hasty J, Johnson TL. 2016. Posttranscriptional Regulation of Gcr1 Expression and Activity Is Crucial for Metabolic Adjustment in Response to Glucose Availability. Mol Cell 62:346–358. doi:10.1016/j.molcel.2016.04.012

Irmisch A, Ampatzidou E, Mizuno K, O’Connell MJ, Murray JM. 2009. Smc5/6 maintains stalled replication forks in a recombination-competent conformation. EMBO J. doi:10.1038/emboj.2008.273

Jeppsson K, Carlborg KK, Nakato R, Berta DG, Lilienthal I, Kanno T, Lindqvist A, Brink MC, Dantuma NP, Katou Y, Shirahige K, Sjögren C. 2014. The Chromosomal Association of the Smc5/6 Complex Depends on Cohesion and Predicts the Level of Sister Chromatid Entanglement. PLoS Genet 10. doi:10.1371/journal.pgen.1004680

Johnson ES, Schwienhorst I, Dohmen RJ, Blobel G. 1997. The ubiquitin-like protein Smt3p is activated for conjugation to other proteins by an Aos1p/Uba2p heterodimer. EMBO J 16:5509–5519. doi:10.1093/emboj/16.18.5509

Jorgensen P, Tyers M. 2004. How cells coordinate growth and division. Curr Biol 14:1014– 1027. doi:10.1016/j.cub.2004.11.027

Joshua IM, Höfken T. 2017. From lipid homeostasis to differentiation: Old and new functions of the zinc cluster proteins Ecm22, Upc2, Sut1 and Sut2. Int J Mol Sci 18:1–17. doi:10.3390/ijms18040772

Kato K, Yamamoto Y, Izawa S. 2011. Severe ethanol stress induces assembly of stress granules in Saccharomyces cerevisiae. Yeast 28:339–347. doi:10.1002/yea.1842

Kato S, Yoshida M, Izawa S. 2019. Btn2 is involved in the clearance of denatured proteins caused by severe ethanol stress in Saccharomyces cerevisiae. FEMS Yeast Res 19:1–8. doi:10.1093/femsyr/foz079

Keller A, Nesvizhskii AI, Kolker E, Aebersold R. 2002. Empirical statistical model to estimate the accuracy of peptide identifications made by MS/MS and database search. Anal Chem 74:5383–5392. doi:10.1021/ac025747h

Lewicki MC, Srikumar T, Johnson E, Raught B. 2015. The S. cerevisiae SUMO stress response is a conjugation-deconjugation cycle that targets the transcription machinery. J Proteomics 118. doi:10.1016/j.jprot.2014.11.012

Lewis JA, Elkon IM, McGee MA, Higbee AJ, Gasch AP. 2010. Exploiting natural variation in Saccharomyces cerevisiae to identify genes for increased ethanol resistance. Genetics 186:1197–1205. doi:10.1534/genetics.110.121871

Liang J, Li BZ, Tan AP, Kolodner RD, Putnam CD, Zhou H. 2018. SUMO E3 ligase Mms21 prevents spontaneous DNA damage induced genome rearrangements. PLoS Genet 14. doi:10.1371/journal.pgen.1007250

Liu G, Bergenholm D, Nielsena J. 2016. Genome-wide mapping of binding sites reveals multiple biological functions of the transcription factor Cst6p in Saccharomyces cerevisiae. MBio 7:1–10. doi:10.1128/mBio.00559-16

Ma Y, Kanakousaki K, Buttitta L. 2015. How the cell cycle impacts chromatin architecture and influences cell fate. Front Genet 5:1–18. doi:10.3389/fgene.2015.00019

Mayhew D, Mitra RD. 2014. Transcription factor regulation and chromosome dynamics during pseudohyphal growth. Mol Biol Cell 25:2669–2676. doi:10.1091/mbc.E14-04-0871

McIsaac RS, Petti AA, Bussemaker HJ, Botstein D. 2012. Perturbation-based analysis and modeling of combinatorial regulation in the yeast sulfur assimilation pathway. Mol Biol Cell 23:2993–3008. doi:10.1091/mbc.E12-03-0232

Menolfi D, Delamarre A, Lengronne A, Pasero P, Branzei D. 2015. Essential Roles of the Smc5/6 Complex in Replication through Natural Pausing Sites and Endogenous DNA Damage Tolerance. Mol Cell 60:835–846. doi:10.1016/j.molcel.2015.10.023

Miller MJ, Scalf M, Rytz TC, Hubler SL, Smith LM, Vierstra RD. 2013. Quantitative proteomics reveals factors regulating RNA biology as dynamic targets of stress-induced SUMOylation in arabidopsis. Mol Cell Proteomics 12:449–463. doi:10.1074/mcp.M112.025056

Mohd Azhar SH, Abdulla R, Jambo SA, Marbawi H, Gansau JA, Mohd Faik AA, Rodrigues KF. 2017. Yeasts in sustainable bioethanol production: A review. Biochem Biophys Reports 10:52–61. doi:10.1016/j.bbrep.2017.03.003

Munk S, Sigurðsson JO, Xiao Z, Batth TS, Franciosa G, von Stechow L, Lopez-Contreras AJ, Vertegaal ACO, Olsen JV. 2017. Proteomics Reveals Global Regulation of Protein SUMOylation by ATM and ATR Kinases during Replication Stress. Cell Rep 21:546–558. doi:10.1016/j.celrep.2017.09.059

Nacerddine K, Lehembre F, Bhaumik M, Artus J, Cohen-Tannoudji M, Babinet C, Pandolfi PP, Dejean A. 2005. The SUMO pathway is essential for nuclear integrity and chromosome segregation in mice. Dev Cell 9:769–779. doi:10.1016/j.devcel.2005.10.007

Nadel CM, Mackie TD, Gardner RG. 2019. Osmolyte accumulation regulates the SUMOylation and inclusion dynamics of the prionogenic Cyc8-Tup1 transcription corepressor. PLoS Genet 15:1–23. doi:10.1371/journal.pgen.1008115

Nishimura K, Fukagawa T, Takisawa H, Kakimoto T, Kanemaki M. 2009. An auxin-based degron system for the rapid depletion of proteins in nonplant cells. Nat Methods 6:917– 922. doi:10.1038/nmeth.1401

Niskanen EA, Palvimo JJ. 2017. Chromatin SUMOylation in heat stress: To protect, pause and organise?: SUMO stress response on chromatin. BioEssays. doi:10.1002/bies.201600263

Oeser ML, Amen T, Nadel CM, Bradley AI, Reed BJ, Jones RD, Gopalan J, Kaganovich D, Gardner RG. 2016. Dynamic Sumoylation of a Conserved Transcription Corepressor Prevents Persistent Inclusion Formation during Hyperosmotic Stress. PLoS Genet 12. doi:10.1371/journal.pgen.1005809

Okuma T, Honda R, Ichikawa G, Tsumagari N, Yasuda H. 1999. In vitro SUMO-1 modification requires two enzymatic steps, E1 and E2. Biochem Biophys Res Commun 254:693–698. doi:10.1006/bbrc.1998.9995

Parapouli M, Vasileiadis A, Afendra AS, Hatziloukas E. 2020. Saccharomyces cerevisiae and its industrial applications, AIMS Microbiology. doi:10.3934/microbiol.2020001

Pohlers S, Martin R, Hellwig D, Kniemeyer O, Saluz HP, Dijck P Van, Ernst JF, Brakhage A, Kurzai O. 2017. Lipid Signaling via Pkh1/2 Regulates Fungal CO2 Sensing through the Kinase Sch9. MBio 8:1–15.

Qu Y, Jiang J, Liu X, Wei P, Yang X, Tang C. 2019. Cell Cycle Inhibitor Whi5 Records Environmental Information to Coordinate Growth and Division in Yeast. Cell Rep 29:987-994.e5. doi:10.1016/j.celrep.2019.09.030

Richardson LA, Reed BJ, Charette JM, Freed EF, Fredrickson EK, Locke MN, Baserga SJ, Gardner RG. 2012. A Conserved Deubiquitinating Enzyme Controls Cell Growth by Regulating RNA Polymerase I Stability. Cell Rep 2:372–385. doi:10.1016/j.celrep.2012.07.009

Sanchez Y, Desany BA, Jones WJ, Liu Q, Wang B, Elledge SJ. 1996. Regulation of RAD53 by the ATM-like kinases MEC1 and TEL1 in yeast cell cycle checkpoint pathways. Science (80-). doi:10.1126/science.271.5247.357

Seeber A, Gasser SM. 2017. Chromatin organization and dynamics in double-strand break repair. Curr Opin Genet Dev 43:9–16. doi:10.1016/j.gde.2016.10.005

Stanley D, Bandara A, Fraser S, Chambers PJ, Stanley GA. 2010. The ethanol stress response and ethanol tolerance of Saccharomyces cerevisiae. J Appl Microbiol 109:13– 24. doi:10.1111/j.1365-2672.2009.04657.x

Steensels J, Verstrepen KJ. 2014. Taming Wild Yeast: Potential of Conventional and Nonconventional Yeasts in Industrial Fermentations. Annu Rev Microbiol 68:61–80. doi:10.1146/annurev-micro-091213-113025

Stielow B, Sapetschnig A, Krüger I, Kunert N, Brehm A, Boutros M, Suske G. 2008. Identification of SUMO-Dependent Chromatin-Associated Transcriptional Repression Components by a Genome-wide RNAi Screen. Mol Cell 29:742–754. doi:10.1016/j.molcel.2007.12.032

Talamillo A, Barroso-Gomila O, Giordano I, Ajuria L, Grillo M, Mayor U, Barrio R. 2020. The role of SUMOylation during development. Biochem Soc Trans 48:463–478. doi:10.1042/BST20190390

Tanaka K, Nishide J, Okazaki K, Kato H, Niwa O, Nakagawa T, Matsuda H, Kawamukai M, Murakami Y. 1999. Characterization of a Fission Yeast SUMO-1 Homologue, Pmt3p, Required for Multiple Nuclear Events, Including the Control of Telomere Length and Chromosome Segregation. Mol Cell Biol 19:8660–8672. doi:10.1128/mcb.19.12.8660

Tempé D, Piechaczyk M, Bossis G. 2008. SUMO under stressBiochemical Society Transactions. pp. 874–878. doi:10.1042/BST0360874

Toone WM, Jones N. 1998. Stress-activated signalling pathways in yeast. Genes to Cells 3:485–498. doi:10.1046/j.1365-2443.1998.00211.x

Tóth ME, Vígh L, Sántha M. 2014. Alcohol stress, membranes, and chaperones. Cell Stress Chaperones. doi:10.1007/s12192-013-0472-5

Tsuyama T, Inou K, Seki M, Seki T, Kumata Y, Kobayashi T, Kimura K, Hanaoka F, Enomoto T, Tada S. 2006. Chromatin loading of Smc5/6 is induced by DNA replication but not by DNA double-strand breaks. Biochem Biophys Res Commun. doi:10.1016/j.bbrc.2006.10.133

Voordeckers K, Colding C, Grasso L, Pardo B, Hoes L, Kominek J, Gielens K, Dekoster K, Gordon J, Van der Zande E, Bircham P, Swings T, Michiels J, Van Loo P, Nuyts S, Pasero P, Lisby M, Verstrepen KJ. 2020. Ethanol exposure increases mutation rate through error-prone polymerases. Nat Commun 11. doi:10.1038/s41467-020-17447-3

Wu CS, Ouyang J, Mori E, Nguyen HD, Maréchal A, Hallet A, Chen DJ, Zou L. 2014. SUMOylation of ATRIP potentiates DNA damage signaling by boosting multiple protein interactions in the ATR pathway. Genes Dev 28:1472–1484. doi:10.1101/gad.238535.114

Yang KM, Lee NR, Woo JM, Choi W, Zimmermann M, Blank LM, Park JB. 2012. Ethanol reduces mitochondrial membrane integrity and thereby impacts carbon metabolism of Saccharomyces cerevisiae. FEMS Yeast Res 12:675–684. doi:10.1111/j.1567-1364.2012.00818.x

Zapatka M, Pociño-Merino I, Heluani-Gahete H, Bermúdez-López M, Tarrés M, Ibars E, Solé-Soler R, Gutiérrez-Escribano P, Apostolova S, Casas C, Aragon L, Wellinger R, Colomina N, Torres-Rosell J. 2019. Sumoylation of Smc5 Promotes Error-free Bypass at Damaged Replication Forks. Cell Rep. doi:10.1016/j.celrep.2019.10.123

Zhao X. 2018. SUMO-Mediated Regulation of Nuclear Functions and Signaling Processes. Mol Cell. doi:10.1016/j.molcel.2018.07.027

Zhou W, Ryan JJ, Zhou H. 2004. Global analyses of sumoylated proteins in Saccharomyces cerevisiae. Induction of protein sumoylation by cellular stresses. J Biol Chem. doi:10.1074/jbc.M404173200

